# Preferential selection and contribution of non-structural protein 1 (NS1) to the efficient transmission of the panzootic avian influenza H5N8 2.3.4.4 clades A and B viruses in chickens and ducks

**DOI:** 10.1101/2021.03.14.435362

**Authors:** Claudia Blaurock, Angele Breithaupt, David Scheibner, Ola Bagato, Axel Karger, Thomas C. Mettenleiter, Elsayed M. Abdelwhab

## Abstract

Highly pathogenic avian influenza viruses H5N8 clade 2.3.4.4 caused outbreaks in poultry at an unprecedented global scale. The virus was spread by wild birds in Asia in two waves: clade-2.3.4.4A in 2014/2015 and clade-2.3.4.4B since 2016 up to today. Both clades were highly virulent in chickens, but only clade-B viruses exhibited high virulence in ducks. Viral factors which contribute to virulence and transmission of these panzootic H5N8 2.3.4.4 viruses are largely unknown. The NS1 protein, typically composed of 230 amino acids (aa), is a multifunctional protein which is also a pathogenicity factor. Here, we studied the evolutionary trajectory of H5N8 NS1 proteins from 2013 to 2019 and their role in the fitness of H5N8 viruses in chickens and ducks. Sequence analysis and *in-vitro* experiments indicated that clade-2.3.4.4A and clade-2.3.4.4B viruses have a preference for NS1 of 237-aa and 217-aa, respectively over NS1 of 230-aa. NS217 was exclusively seen in domestic and wild birds in Europe. The extension of the NS1 C-terminus of clade-B virus reduced virus transmission and replication in chickens and ducks and partially impaired the systemic tropism to the endothelium in ducks. Conversely, lower impact on fitness of clade-A virus was observed. Remarkably, the NS1 of clade-A and clade-B, regardless of length, was efficient to block interferon induction in infected chickens and changes in the NS1 C-terminus reduced the efficiency for interferon antagonism. Together, the NS1 C-terminus contributes to the efficient transmission and high fitness of H5N8 viruses in chickens and ducks.

**Importance:** The panzootic H5N8 highly pathogenic avian influenza viruses of clade 2.3.4.4A and 2.3.4.4B devastated poultry industry globally. Clade 2.3.4.4A was predominant in 2014/2015 while clade 2.3.4.4B was widely spread in 2016/2017. Both clades exhibited different pathotypes in ducks. Virus factors contributing to virulence and transmission are largely unknown. The NS1 protein is typically composed of 230 amino-acids (aa) and is an essential interferon (IFN) antagonist. Here, we found that the NS1 protein of clade 2.3.4.4A preferentially evolved toward long NS1 with 237-aa, while clade 2.3.4.4B evolved toward shorter NS1 with 217-aa (exclusively found in Europe) due stop-codons in the C-terminus (CTE). We showed that the NS1 CTE of H5N8 is required for efficient virus replication, transmission and endotheliotropism in ducks. In chickens, H5N8 NS1 evolved toward higher efficiency to block IFN-response. These findings may explain the preferential pattern for short NS1 and high fitness of the panzootic H5N8 in birds.

**Subject category:** Animal, RNA Viruses

## Introduction

Avian influenza viruses (AIV) infect a wide range of birds and mammals exhibiting high pathogenic (HP) or low pathogenic (LP) phenotypes. AIV belongs to genus Influenza A Virus (IAV) in the family *Orthomyxoviridae* with an RNA genome of eight gene segments (segments 1 to 8) which encode more than ten structural and non-structural proteins (1). The non-structural protein-1 (NS1), encoded by segment 8 encompassing 890 nucleotides, occurs in two distinct alleles (phylogroups) in birds, where allele A is more common than allele B (2-4). NS1 typically encompasses 230 amino acids (aa) arranged in two domains: the RNA binding domain (RBD) from aa 1 to 73 and an effector domain (ED) from aa 88-230 which are connected by a linker (residues 74-87) (5). It is present as a homodimer. Upon infection of cells, the interactions of NS1 RBD with many viral and host RNA species prevents the activation of host cellular sensors and suppresses cellular gene expression. Likewise, the ED interacts with a plethora of host proteins to, among other functions, antagonize host immune response (5, 6).

Although, NS1 has a typical length of 230 aa, several influenza viruses (e.g. H5Nx) have a shorter NS1 due to a 5-aa deletion in the linker region (80-84 aa) (5) or a deletion in the C-terminus (ΔCTE) of the ED. We and others showed that there are up to 13 different forms of the ΔCTE in AIV (4, 7, 8). The most common ΔCTE form (88%) was NS with 217-aa lacking aa 218-230, while the extension in CTE to 237 aa was rarely observed (n= 112/13026, 0.9%) (8). Generally, in poultry the impact of NS1 CTE variations on virulence of AIV is controversial and the contribution to virus transmission or tropism was rarely studied. Species-specific variations in the PDZ domain (aa 227-230) of NS1 affected LPAIV H7N1 and HPAIV H5N1 replication in duck cell cultures but not in chicken embryo fibroblasts (9, 10). *In-vivo*, no significant difference in virulence of HPAIV H5N1 in chickens was observed due to variation of the PDZ domain (10) and it was dispensable for virus replication in chickens and ducks (11). Conversely, a deletion of the PDZ domain in the HPAIV H7N1 NS1 was advantageous for virus replication in chicken embryo fibroblasts (8, 12) and in infected chickens (8) but did not significantly affect the high virulence of the virus in infected chickens (8). In another study, extension of 217-aa NS1 to 230 or 237 aa did not have a significant impact on LPAIV H9N2 replication in avian cells, but increased virus replication and transmission in chickens without significant impact on virulence (13). No information is available on the impact of CTE elongation on HPAIV fitness, particularly transmissibility, in ducks.

The HPAIV Goose/Guangdong (GsGd96) H5N1 first reported in Hong Kong in 1996/1997, with full NS1 CTE, devastated poultry in more than 60 countries (14) and 455 out of 861 (53%) infected humans died (15). Since then, the virus evolved into tens of clades and subclades and underwent several reassortment events. In recent years, H5N8 clade 2.3.4.4 spread globally in an unprecedented panzootic (14). In 2013-2015, H5N8 clade 2.3.4.4A (designated H5N8-A) was transported by wild birds from Asia to birds and poultry in Europe and North America (16, 17). In 2016, the second wave of H5N8 caused by clade 2.3.4.4B (designated H5N8-B) resulted in devastating outbreaks in poultry in several countries in Asia, Europe, Africa and the Middle East (18-20). H5N8-A viruses were avirulent in Pekin ducks, while H5N8-B viruses were highly virulent (21). We have recently shown that NS1 is a major virulence determinant of an H5N8-B virus in ducks (Scheibner et al. in preparation) and swapping NS gene segment of H5N8-B and LPAIV H9N2 reduced virus transmission in chickens (22). Little is known about the global evolution of NS1 gene of clade 2.3.4.4 viruses overtime and the role of CTE on virus fitness in gallinaceous birds and waterfowls. In this study, we analysed all sequence of NS1 of clade 2.3.4.4 H5Nx viruses and studied the impact of NS1 CTE deletion and extension on replication, virulence and transmission of recent HPAIV H5N8-A and H5N8-B in chickens and ducks.

## Materials and Methods

### Sequence analysis

A total of 8185 complete NS1 protein sequences were retrieved from GISAID (n= 2872) and Influenza Virus Resources (n= 5313) available publicly from 1996 to 2019. Sequences were edited manually (e.g. to remove the laboratory-viruses and viruses with ambiguous sequences or nomenclature, etc.), aligned using MAFFT and further edited using Geneious version 2019.2.3. Furthermore, statistical programming language R (23) was used to analyse the prevalence of sequences of H5N8 viruses from 2013 to 2019 and the R package ggplot2 (24) to construct graphs. MrBayes was used to analyse the phylogenetic relatedness after selection of bestfit Model in Topali v2 software (25) and further edited using Dendroscope and Inkscape free software.

### Viruses and cell lines

A/turkey/Germany-MV/AR2487/2014 (H5N8) carrying NS with 237-aa (designated hereafter A_NS237) and A/tufted duck/Germany/8444/2016 (H5N8) carrying NS with 217-aa (designated hereafter B_NS217) were kindly provided by Timm C. Harder, the head of reference laboratory for avian influenza virus, Friedrich-Loeffler-Institut (FLI), Greifswald Insel-Riems, Germany. The HA of both viruses belonged to clade 2.3.4.4A and 2.3.4.4B, respectively. Madin-Darby canine kidney type II cells (MDCKII) and human embryonic kidney cells (HEK293T) were provided by the Cell Culture Collection in Veterinary Medicine of the FLI. Primary chicken embryo kidney (CEK) and duck embryo kidney (DEK) cells were prepared according to the standard protocol and as described below (26).

### Construction of recombinant wild and mutant viruses

All viruses were handled in biosafety level 3 (BSL3) facilities at the FLI. Recombinant B_NS217 virus was generated in a previous study (22). To generate a recombinant A_NS237 virus, viral RNA was extracted from allantoic fluid using Trizol reagent (Thermo Fischer, Germany) and Qiagen RNeasy Kit (Qiagen, Germany) following the manufacturers’ guidelines. cDNA was transcribed using reverse primer targeting the conserved termini of influenza gene segments and the Omniscript Reverse Transcription Kit (Qiagen, Germany). Each gene segment was amplified using segment-specific primers and extracted from 1% gel slices using Qiagen Gel Extraction Kit (Qiagen, Germany). Purified gene segments were cloned into the plasmid pHW*Sccd*B as previously done (27). Plasmids containing different gene segments were extracted from transformed *E. coli* XL1-Blue™ or SURE2™ Supercompetent Cells (Agilent, Germany) using QIAGEN Plasmid Mini and Midi Kit (Qiagen, Germany) and subjected for Sanger sequencing (Eurofins, Germany). The NS gene segments of A_NS237 and B_NS217 were modified using QuikChange II Site Directed Mutagenesis Kit according to the instruction manual (Agilent, Germany). The sequence of mutagenesis primers is available upon request.

HEK293T/MDCKII co-culture were transfected with 1 µg plasmids of each gene segment (27) in a mixture of Opti-MEM containing GlutaMAX and Lipofectamine 2000 (Fischer Scientific, Germany). After 2 days, 9-11 day-old specific pathogen free (SPF) embryonated chicken eggs (ECE) (VALO BioMedia GmbH, Germany) were inoculated via the allantoic sac with the supernatant of transfected cells (28). Eggs were checked daily for embryo mortality and the hemagglutination activity in the allantoic fluid was tested against 1% chicken erythrocytes following the standard protocol. Allantoic fluids with titre higher than 16 hemagglutination units were tested for bacterial contamination using Columbia sheep blood agar (Thermo Fisher, Germany). Virus stocks were dispensed in 2 mL-cryotubes and stored at −80°C until further use.

### Virus titration

Virus titration in this study was done using standard plaque assay. Briefly, viruses were serially diluted in ten-fold dilutions in PBS and were added to MDCKII cells in 12-well plates for 1 h at 37°C/5% CO_2_. Cells were washed with phosphate buffer saline (PBS, pH= 7.4). Semisolid agar (Bacto™ Agar, BD, France) and minimal essential medium (MEM) containing 4% bovine serum albumin (BSA) (MP Biomedicals, USA) were mixed to equal parts and was added to each well. After three days at 37°C, cells were fixed by 4% formaldehyde containing 0.1% crystal violet for 24 h and plaques were counted. Viral titres were expressed as plaque forming units per ml (PFU/ml).

### Preferential selection of NS segment in vitro

To determine the preferential selection of the authentic NS segment (NS237 in clade-A or NS217 in clade-B) over NS230, HEK293T cells were co-transfected with 9 plasmids in 4 different settings. Cells were co-transfected with plasmids containing all gene segments of clade-A A_NS217 virus in addition to clade-A NS segments with NS230 or NS237, each in two independent experiments. Similarly, cells were co-transfected with plasmids containing all gene segments of clade-B B_NS237 virus in addition to clade-B NS segments with NS230 or NS217. Transfection was done as described above using OptiMEM and Lipofectamine. Two days after transfection, 2 eggs per transfection were inoculated with the transfected-cell supernatant and daily checked for embryo mortality. The allantoic fluid was harvested and plaque assay was conducted. A total of 70 plaques were randomly selected and RNA was extracted using NucleoSpin 96 Virus Core Kit (Macherey & Nagel GmbH, Germany). cDNA Synthesis was performed with OneStep RT-PCR Kit (Qiagen, Germany) and NS specific primers. Purified gel products were subjected for Sanger sequencing (Eurofins, Germany). Prevalence of different NS variants was analysed using Geneious version 2019.2.3.

### Replication kinetics in primary chicken embryo kidney (CEK) and duck embryo kidney (DEK) cells

CEK cells were prepared from 18-day-old SPF ECE and DEK cells were prepared from 23-day-old duck eggs. Briefly, embryos were decapitated using sterile scissors and the kidney was removed in sterile petri dish. Cell suspensions were prepared by trypsinization and mechanically by sterile scissors in presence of MEM containing 10% foetal calf serum (FCS) (Biowest, Germany). Trypsinized cells were collected in a flask containing MEM and subjected for stirring using magnetic bar at 200 rpm at room temperature (rt) for 25 minutes. The suspension was purified by decanting the whole amount in gauze pads in a beaker followed by centrifugation for 5 min at 1200 rpm. The pellet was suspended in MEM containing 10% FCS, penicillin-streptomycin (1:100) (Thermo Fisher Scientific, USA) and amphotericin B (1:1000) (Biowest, Germany). Finally, ∼500,000 cells per well were distributed in 12-well plates. After 24 h, semi-confluent cells were infected at a multiplicity of infection (MOI) of 1 PFU per 1000 cells in MEM and left for 1 h at 37°C/5% CO_2_. Thereafter, the inoculum was removed and extracellular virions were inactivated by citric acid buffer (pH= 3) for two minutes and the cells were washed twice with PBS. MEM with 0.2% BSA was added to each well. All plates were incubated at 37°C/5% CO_2_. At indicated hours post infection (hpi), cells were collected and kept at −80°C until use. The assay was run in duplicates for each virus and repeated twice. Virus titres were determined by plaque assay in MDCKII as described above. The results were expressed as mean and standard deviation of all replicates.

### Protein expression in avian cells using Western Blot

To study the impact of CTE on expression of NS1, CEK cells were infected at an MOI of 0.1 in duplicates for 1 h at 37°C and 40 °C. Cells were treated with citrate buffer saline for 2 minutes then washed twice with PBS before adding MEM containing 0.2 % BSA and further incubation for 2, 4, 8 and 24 hours. At the indicated time points, cells were collected and subjected for two rounds of centrifugation for 10 min at 13,000 rpm and washing by PBS. Cell pellets were dissolved in Laemmli buffer (SERVA, Germany) and PBS at ratio 1:1 and heated for 5 min at 95° C. Proteins were separated in standard procedures using 12% polyacrylamide gel (SDS-PAGE) and transferred to polyvinylidene fluoride (PVDF) membranes (GE Healthcare Life science, Germany) using TransBlot semi dry transfer cell (Bio-Rad, Germany). After saturation of non-specific protein binding using 5% low-fat milk diluted in TBS-T at rt for one hour, the membranes were incubated overnight under constant shaking at 4°C with the primary polyclonal rabbit anti-NS1 antibody. Furthermore, β-actin as well as NP were detected using monoclonal anti-β-actin (Sigma-Aldrich, USA) as well as polyclonal-anti-NP rabbit antibodies, respectively. After washing the membrane three times with 1xTBS-T (each step 5 min), secondary peroxidase-conjugated rabbit and mouse IgG at a dilution of 1:20000 in TBS-T were added to the membrane for 1 h at rt. Thereafter, the membranes were washed 3 times with 1 x TBS-T and antibody binding was detected in a BioRad Versa Doc System with the Quantity One software (BioRad, Germany) by chemiluminescence using the Clarity Western ECL Substrate (BioRad, Germany).

### Animal experiments

#### Ethic statement

All experiments in this study were carried out according to the German Regulations for Animal Welfare after obtaining the necessary approval from the authorized ethics committee of the State Office of Agriculture, Food Safety, and Fishery in Mecklenburg – Western Pomerania (LALLF M-V) under permission number: 7221.3-1-060/17 and approval of the commissioner for animal welfare at the FLI representing the Institutional Animal Care and Use Committee (IACUC). All animal experiments were conducted in the BSL3 animal facilities at the FLI.

#### Experimental design

The impact of CTE on virulence and transmission was assessed in White Leghorn chickens and Pekin ducks. One-day-old SPF chicks hatched at the FLI, Celle, Germany and one-day old Pekin ducklings (Duck-Tec Brueterei, Germany) were used for this study. Ducks were tested to exclude bacterial (e.g. Salmonella spp.) and viral (i.e. influenza) infections. Two days before challenge, 15 chickens (6-week old) and ducks (2-week old) were allocated into six groups in separate experimental rooms. Ten birds were challenged with indicated viruses by oculonasal (ON) inoculation and 1 day-post-inoculation (dpi), 5 naïve birds were added to each inoculated group to assess bird-to-bird transmission. All birds were observed daily for morbidity and mortality for 10 days. Clinical scoring was adopted as previously done (8): 0 for clinically healthy birds, 1 for moribund birds showing one clinical sign including respiratory disorders, cyanosis, nervous signs or diarrhoea and 2 for moribund birds showing two or more clinical signs and 3 for dead birds. Severely moribund birds which were not able to eat or drink were humanely killed using isoflurane (CP Pharma, Germany) and were given score 3 at the next day. Furthermore, the pathogenicity index (PI) for each virus was calculated as the mean value for daily-scores of all birds in 10 days divided by 10. The mean time to death (MTD) was calculated by multiplying the number of birds that died per day as day of death divided by the total number of dead birds in each group.

#### Virus excretion

To determine the impact of CTE on virus excretion in inoculated and sentinel chickens and ducks, oropharyngeal (OP) and cloacal (CL) swabs were obtained from surviving or freshly dead birds at 2 and 4 dpi in MEM containing 0.2% BSA and antibiotics. Extraction of viral RNA from swab media was done using Viral RNA/DNA Isolation Kit (NucleoMagVET) (Macherey & Nagel GmbH, Germany) in a KingFisher Flex Purification System (Thermo Fisher Scientific, USA). Partial amplification of AIV-M gene was done using SuperScript III One-Step RT-PCR System with Platinum Taq DNA Polymerase (Invitrogen, Germany) and generic real-time reverse-transcription PCR (RT-qPCR) as previously published (29) in AriaMx real-time cycler (Agilent, Germany). Standard curves generated by serial dilutions of B_NS217 (10 to 100000 pfu) were used in each plate. To semi-quantify the viral RNA, plotting of the Ct-value and the corresponding dilution in the standard curve was automatically done. Results are shown as mean and standard deviation of positive birds.

#### Seroconversion

At the end of the experiment, blood samples were collected from the jugular vein in all surviving birds after deep anaesthesia. Serum samples were collected after 24 h at 4°C and centrifugation. Anti-AIV NP was detected using ID screen Influenza A Antibody Competition Multispecies ELISA Kit (IDvet, France). Plates were read in Infinite 200 PRO reader (Tecan Trading AG, Switzerland).

#### Histopathology and immunohistochemistry

To determine the impact of CTE on microscopic lesions and distribution of AIV in different tissues, samples were collected from freshly dead or slaughtered inoculated birds (n=3 per group) at 2 dpi. Samples from beak, trachea, lungs, brain, heart, pancreas, liver, kidney, spleen, proventriculus, gizzard, duodenum, cecum, cecal tonsils, bursa of Fabricius and thymus were collected in 4% phosphate-buffered neutral formaldehyde. The tissues were embedded in paraffin, cut at 3 µm thickness and stained with haematoxylin and eosin (HE) for light microscopical examination (blinded study). Scoring of microscopic lesions was performed on a necrosis scale 0 to 3+: 0 = no lesion, 1+ = mild, 2+ = moderate, 3+ = severe. Furthermore, viral antigen detection was performed using the avidin-biotin-complex (ABC) in immunohistochemistry (IHC) method (30). The M1 protein of influenza A virus was detected by ATCC HB-64 monoclonal antibody diluted 1:200 in Tris-buffered saline, pH 7.6 at 4°C overnight and goat anti-mouse IgG (#BA 9200, Vector Laboratories, Burlingame, CA) diluted 1:200. Bright red signals were generated with an ABC Kit (Vectastain Elite #PK6100, Vector Laboratories) and the AEC substrate 3-amino-9-ethylcarbazole (Dako, Carpinteria, CA). Moreover, sections were counterstained with Mayer’s haematoxylin. Positive control tissue samples and a nonrelated control antibody were included in each run. Scoring was done on a scale 0 to 3+: 0 = negative, 1+ = focal to oligofocal, 2+ = multifocal, 3+ = coalescing foci or diffuse labeling.

#### Detection of cytokines in chickens and ducks

To get an insight into the impact of CTE on immune response, interferons were measured by generic RT-qPCR (31, 32). Chickens and ducks (n=3) were inoculated with different viruses in a separate experiment. Lungs and spleen of chickens were collected at 2 dpi, weighed (w/v) and homogenized using TissueLyzer® (Qiagen, Germany). The total RNA was extracted from homogenized tissues using Trizol and RNeasy Kit following the manufacturer’s instructions (Qiagen, Germany). The cDNA was transcribed from 400 ng RNA by the Prime Script 1^st^ Strand cDNA Synthesis Kit (TaKaRa, Germany) and Oligo dT Primer (TaKaRa) in a total 20 µl reaction as recommended by the producers. Quantification of IFN-α, IFN-β and IFN-γ transcripts was done using TaqMan probes for chickens or SYBR GreenER qPCR SuperMIX Universal Kit (Invitrogen) for ducks (31, 32). Normalization was done using 28S rRNA transcripts (33). Results were calculated using the 2^-(ΔΔct) method and expressed as fold change of normalized samples compared to samples obtained from non-infected birds (n=3).

#### Statistics

Data were analysed for statistics using GraphPad Prism software. Non-parametric Kruskal-Wallis (Dunn’s multiple comparsions) test was used for statistical anaysis of replication kinetics. One-way ANOVA with post hoc Tukey’s test was used for statistical analysis of swab and organ samples as well as for the analysis of the fold-change induction of interferon. Survival time was analysed by Log-rank (Mantel-Cox) test. Data were consideres statistically significant at p-value<0.05.

## Results

### Temporal and spatial variations in the NS1 C-terminus of clade 2.3.4.4 H5N8 viruses

To understand the global evolution of NS1 CTE in H5 viruses, we analysed 8185 NS1 protein sequences from H5Nx viruses available in GenBank and GISAID. Our analysis revealed that NS1 exhibits 15 size variants. The parental virus GsGD96 had a typical NS1 of 230-aa with full CTE. From 1997, other NS1 with variable aa lengths including 202, 212, 215, 217, 220, 223, 224, 225, 227, 228, 230, 232, 236, 237 and 238 aa were found (data not shown). The most predominant forms were NS217, NS225, NS230 and NS237. NS225 and NS230 have a complete CTE, but vary due to a deletion of aa 80 to 84. Other variants have deletions or insertions in the CTE. For instance, NS217 and NS237 have a 13-aa deletion or 7-aa insertion in the CTE due to variable stop codons in the NS1 CTE, respectively.

To determine the origin and evolution of the NS segment in H5N8-A and H5N8-B, 1068 H5N8 sequences from 2013 (n=12), 2014/2015 (n= 465) and 2016/2019 (n=591) from Asia and Europe were analysed. The NS1 of both viruses clustered in two phylogroups within the Eurasian lineage along with other contemporary H5N8 viruses from Europe and Asia (Figure 1A). H5N8 sequences had either NS230, NS237 or NS217. In 2013, Asian H5N8 viruses as well as the putative predecessors (H4N2, H11N9, H4N6 and H5N2 (34)) had NS1 with a length of 230-aa (Figure 1B). In 2014/2015, 428/465 (∼92%) sequences were of 237-aa length, while NS230 was reported only in 37/465 (∼8%) (35 from wild birds mainly waterfowl and 2 from domestic chickens and ducks). In 2016/2019, 201/591 (∼34%) and 390/591 (∼66%) sequences had NS230 or NS237, respectively (Figure 1B). NS230 was reported mainly in Asia in wild birds or environment (n= 155/201; ∼77.1%) and domestic birds (n= 44/201; ∼21.9%). Only 2/201 (1%) viruses with NS230 were reported in turkeys in Italy and Poland. NS217 was not reported in Asia (except one sequence from wild birds in Iran), while in Europe NS217 was reported in 214/390 (∼54.9%) from wild birds or environmental samples and 176/390 (∼45.1%) from domestic birds. No single sequence with NS217 was reported in 2013/2015 and no NS237 was found in 2016/2019 in sequences analysed in this study. These results indicate temporal and spatial patterns for NS1 CTE in H5N8 viruses from 2013 to 2019. In Asia, H5N8 with NS230 and NS237 in 2014/2015 were common in wild birds. Conversely, NS217 was reported in wild and domestic birds exclusively in Europe in 2016/2019. These findings suggest a preferential selection for NS237 and NS217 over NS230 in H5N8-A in 2014/2015 and H5N8-B in 2016/2019, respectively.

**Figure 1:**
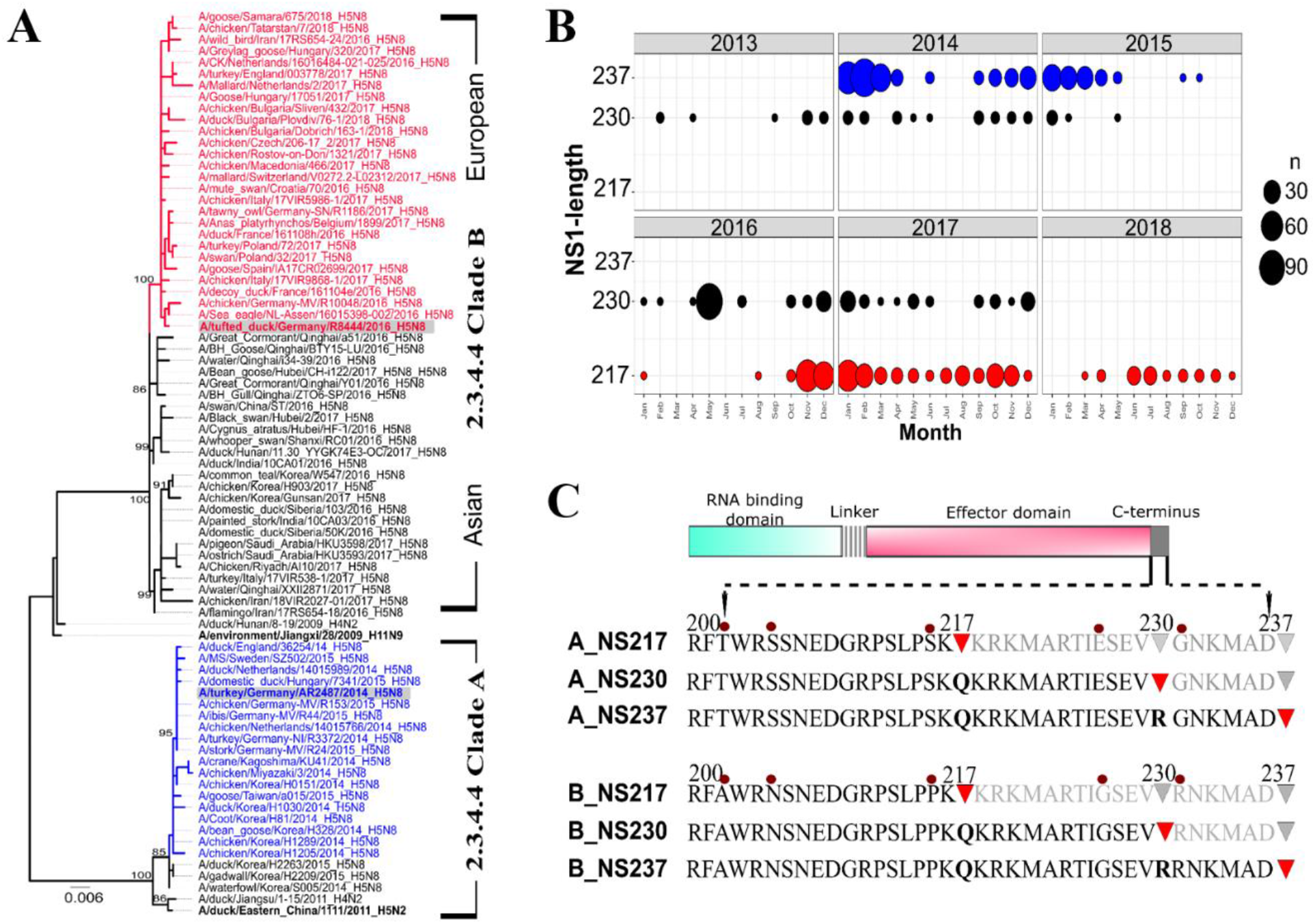
Evolution of NS1 of clade 2.3.4.4A and 2.3.4.4B H5N8 viruses. NS gene sequences were retrieved from GISAID and were aligned using MAFFT. A mid-point rooted phylogenetic tree was generated by MrBayes in Toplai v2. Two independent runs of 1,000,000 replicates and 10% burn-in were used. Shown is the phylogenetic relatedness of NS segments allele A of representative viruses. Clade-A and clade-B viruses used in this study are highlighted in grey. Viruses written in black have NS230, in red have NS217 and in blue have NS237. The putative ancestors are written in bold. Similar topology was obtained using NJ and ML trees (data not shown) (A). R was used to determine the temporal distribution of NS1 length in sequences collected between 2013 to 2018 (B). Recombinant H5N8 clade-A and clade-B 2.3.4.4 viruses were generated carrying different NS1 CTE. Clades A and B wild-type viruses have 237 and 217-aa, respectively. Deep-red circles indicate point mutations in clade-A compared to clade-B. Red triangles indicate stop codons (C).

### Generation of recombinant H5N8-A and H5N8-B viruses with NS217, NS230 and NS237 NS1 proteins

Two H5N8 clade A and clade B viruses were selected for this study. Both viruses were closely related to the contemporary H5N8 viruses (Figure 1A). The NS1 of A_NS237 and B_NS217 viruses share 92.6% nucleotides and 90.8% aa identity with 48 nucleotides and 20 aa differences, respectively. Seven aa differences were found in the RBD (V6M, S7L, H17Y, S48N, D53G, L65V, G70E), 4 in the linker region (N80T, I81V, V84S, T86S) and 9 in the ED (T127N, I129T, D139N, T143A, L166F, I176N, T202A, S205N, S216P). No deletion in aa positions 80-84 was found. Extension of NS1 CTE (clade-A) and truncation (clade-B) was observed (Figure 1C). To study the impact of NS1 CTE on the fitness of H5N8-A and H5N8-B viruses *in vitro* and *in vivo*, recombinant viruses were generated using A_NS237 and B_NS217. We constructed four more recombinant viruses: A_NS230 and A_NS217 were generated from A_NS237 and B_NS230 and B_NS237 were generated from B_NS217. These viruses carry NS1 with different length due to insertion or removal of stop codons in the NS1 CTE (Figure 1C). Sequence of recombinant virus stocks were confirmed by Sanger sequencing and compared to the sequences of the parent viruses. All viruses were propagated in ECE and reached titres ranged from 3.6 × 10^5^ to 2.3 × 10^7^ PFU/ml.

### Preferential selection of H5N8-A and H5N8-B viruses to NS1 with variable C-terminus length

To test whether there is a preference of clade-A and clade-B viruses to certain NS1 variants, we co-transfected HEK293T cells with 8 segments from A_NS237 and plasmids containing A_NS230 or A_NS217. The same experiment was done with 8 segments from B_NS217 and plasmids containing B_NS230 or B_NS237. The experiment was conducted in two independent replicates. After two days, supernatant from transfected cells was inoculated in ECE for two days. Allantoic fluid was serially diluted and titrated in plaque test in MDCKII cells (Figure 2A). Seventy plaques were randomly selected, 37 from clade-A sets and 33 from clade-B sets. RNA was extracted and sequencing of NS segment was generated. Results showed that A_NS237 dominated other clade-A viruses with NS230 and NS217, where 8/16 and 13/21 plaques had NS237 and 8/16 and 8/21 plaques were mixed with A_NS230-NS237 and NS217-NS237, respectively. No single plaque contained A_NS217. Likewise, the B_NS217 dominated other longer NS variants, where 2/11 and 13/22 plaques had NS217, 9/11 and 9/22 plaques were mixed with NS217-NS230 and NS217-NS237 plaques, respectively. No single plaque contained A_NS230 alone (Figure 2A). These results indicate that H5N8 clade-A and clade-B have strain-specific preference for NS1 proteins with long and short C-terminus, respectively.

**Figure 2:**
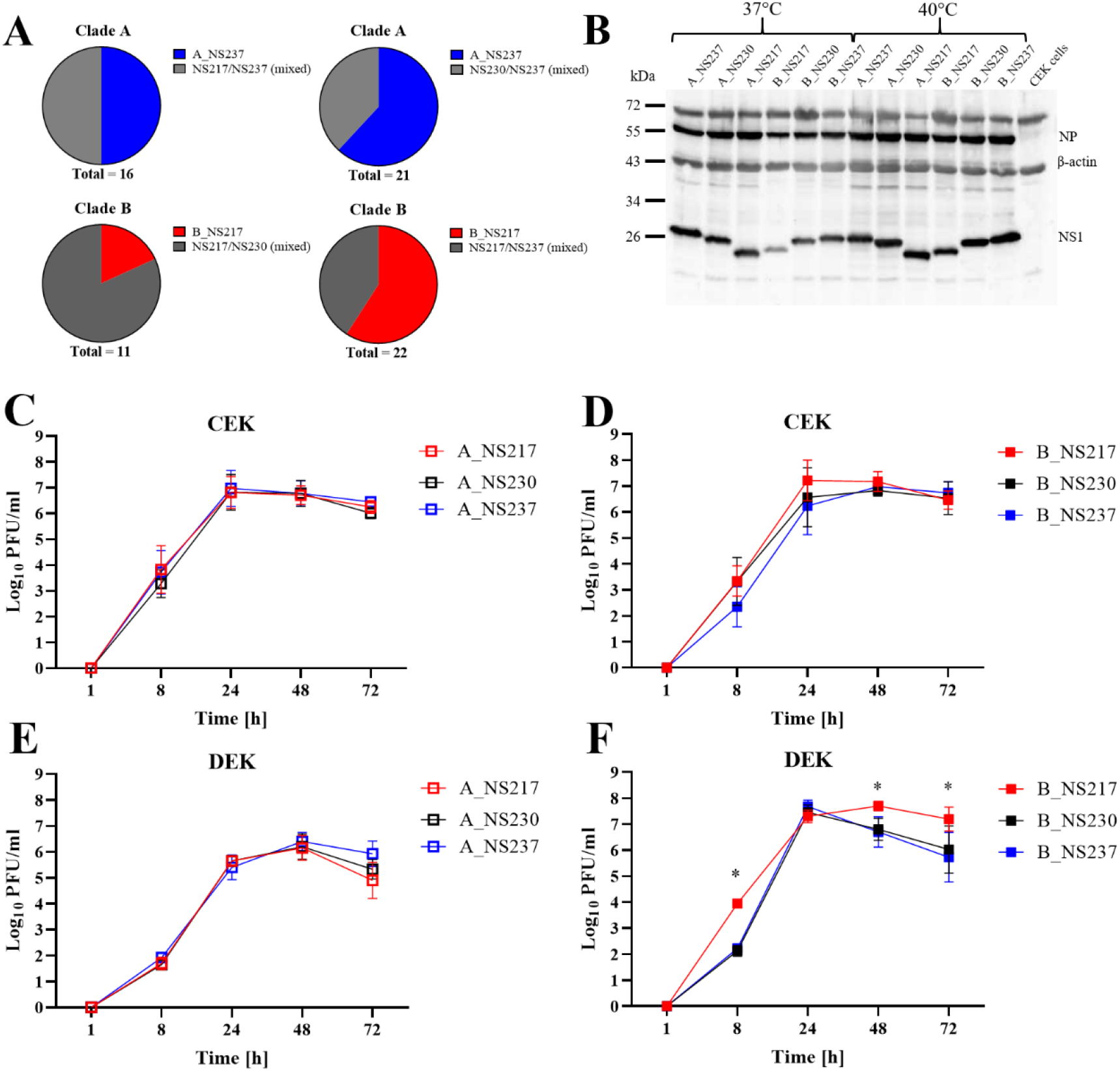
Preferential selection and impact of NS1 CTE on expression and replication of H5N8 viruses in cell culture. Preferential selection of NS1 was studied by co-transfection of cells for 2 days and propagation in ECE. Plaques were randomly selected and NS1 was subjected for Sanger sequencing after RNA extraction and amplification of the NS1. The transfection was run in two separate rounds (A). Expression of NS1 4 hours post infection of CEK cells with an MOI of 0.1. NS1 was detected by rabbit polyclonal NS1 antibodies, NP was detected by polyclonal NP rabbit-antibody and beta-actin with a commercial monoclonal antibody. Similar results were obtained at 8 and 24 hpi (data not shown) (B). Replication of different viruses at MOI of 0.001 at indicated time points in primary embryo kidney cells obtained from chickens (CEK) (C, D) or ducks (DEK) (E, F). Virus titres were determined by plaque assay in MDCKII cells. Statistical significance * p < 0.05.

### Changes in the CTE did not have a significant impact on NS1 expression in avian cells at different temperatures

The expression of NS1 in CEK cells at 2, 4, 8 and 24 hpi at 37°C and 40 °C was studied in duplicates in two independent rounds using Western Blot. At 2 hpi, NS1 was neither detected at 37°C nor at 40 °C (data not shown). At 4, 8 and 24 hpi, the NS1 of all viruses according to the expected molecular weight were detected without significant differences in the amount of NS1 at 37 and 40 °C (Figure 2B). Together, changes in the CTE didn’t affect the NS1 expression in chicken cells at different temperatures.

### Extension of NS1 CTE of H5N8-B reduced virus replication in duck cells, but not in chicken cells

The impact of NS1 CTE on virus replication in primary CEK and DEK cells at 1, 8, 24, 48 and 72 h was studied (Figure 2C-F). No significant differences for the replication of H5N8-A viruses in CEK and DEK and replication of H5N8-B in CEK cells were observed (Figure 2C-E). In DEK cells, B_NS217 replicated to significantly higher titres than B_NS230 and/or B_NS237 at 8, 48 and 72 hpi (p < 0.05) (Figure 2F) and higher than A_NS237 viruses at 24 hpi (p < 0.01). Together, these results indicate that changes in the NS1 CTE did not affect H5N8-A and H5N8-B replication in chicken cells. Conversely, the elongation of the NS1 reduced H5N8-B virus replication in duck cells.

### Elongation of NS1 of H5N8-B virus partially or fully compromised virus transmission to naïve chickens

Virulence of the 6 different viruses was assessed in ten ON-inoculated chickens per group. All birds died within 3 dpi and with the same PI value of 2.7 (Table 1). The MTD was also comparable ranging from 2 to 2.6 days in H5N8-A inoculated groups and 2 to 2.2 days in H5N8-B inoculated groups. All contact chickens in groups inoculated with H5N8-A viruses died within 6 days (5 day-post-contact “dpc”), with MTD of 5.2, 4.8 and 5.4 days in groups co-housed with A_NS237, A_NS230 and A_NS217 inoculated chickens, respectively (Table 1). All chickens co-housed with B_NS217 died within 4 dpi (3 dpc), while 4/5 chickens co-housed with B_NS230 inoculated chickens died within 6 days. None of the contact chickens co-housed with B_NS237 died. Surviving contact chickens in the latter two groups did not seroconvert at 10 dpi (9 dpc) (Table 1). Together, NS1 did not affect virus virulence in chickens. The impact of CTE on chicken-to-chicken transmission is virus dependent. While the elongation of the CTE of H5N8-B compromised virus transmission in chickens, no significant impact on H5N8-A virus was observed.

**Table 1:** Impact of NS1 CTE in chickens after challenge with recombinant H5N8 viruses.

| Clade | Viruses | Inoculated Chickens |  |  | Contact Chickens |  |  |
| --- | --- | --- | --- | --- | --- | --- | --- |
|  |  | Mortality* | Scoring | MTD | Mortality* | MTD | Seroconversion |
| A | A_NS217 | 10/10 | 2.7 | 2.6 | 5/5 | 5.4 | n.a. |
|  | A_NS230 | 10/10 | 2.7 | 2.1 | 5/5 | 4.8 | n.a. |
|  | A_NS237 | 10/10 | 2.7 | 2.0 | 5/5 | 5.2 | n.a. |
| B | B_NS217 | 10/10 | 2.7 | 2.0 | 5/5 | 4.0 | n.a. |
|  | B_NS230 | 10/10 | 2.7 | 2.2 | 4/5 | 5.5 | 0/1 |
|  | B_NS237 | 10/10 | 2.7 | 2.0 | 0/5 | n.a. | 0/5 |

Mortality = number of dead chickens to number of inoculated
MTD= mean time to death
n.a.= not applicable because all birds died

### Changes in the CTE reduced H5N8-B virus oral and cloacal excretion in chickens

Virus excretion at 2 dpi was determined in OP and CL swabs. Viral RNA was detected in all inoculated chickens (Figure 3A, B). RNA levels were comparable in OP and CL swabs in chickens inoculated with H5N8-A viruses. However, B_NS230 was excreted at significantly lower levels than B_NS237 and B_NS217 in the OP and CL swabs (p < 0.04) (Figure 3B). Cloacal excretion of B_NS217 and B_NS237 was higher than in the OP swabs (p < 0.002) (Figure 3B). RNA was not detected in contact chickens in any group. These results indicate that short NS1 CTE is important for the excretion of H5N8-B virus in chickens, but not for H5N8-A.

**Figure 3:**
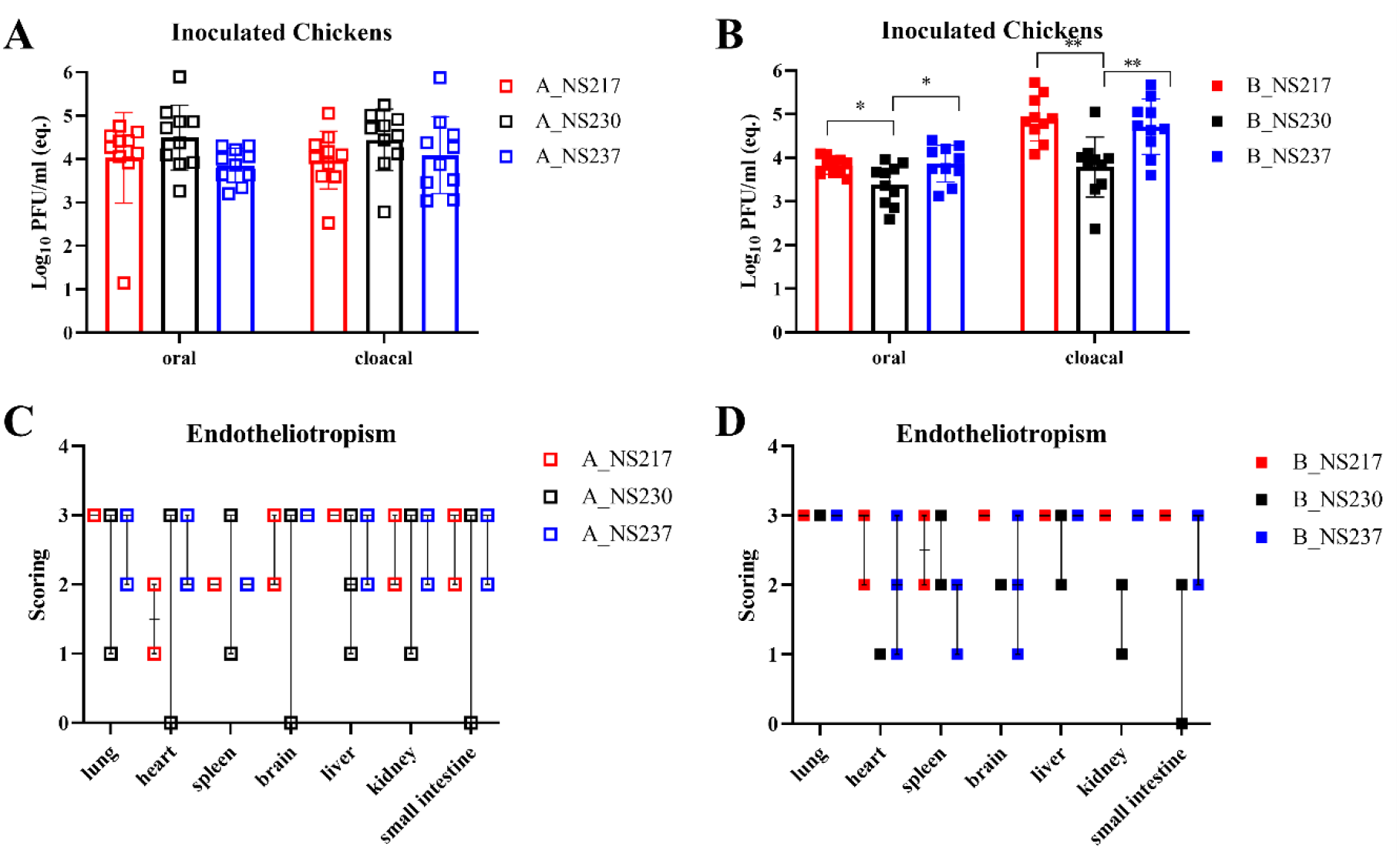
The impact of NS1 on virus excretion and endotheliotropism in chickens. Virus excretion determined in oropharyngeal and cloacal swabs in inoculated chickens 2 dpi using quantitative RT-qPCR was expressed as log10 pfu/ml (eq.) (A, B). Endotheliotropism was determined by IHC. Scores indicate 0 = negative, 1 = focal to oligofocal, 2 = multifocal, 3 = coalescing foci or diffuse labelling, dots represent individual animals, bar = median with interquartile range (C, D). Statistical significance * p < 0.05, ** p< 0.01.

### Changes in the CTE altered H5N8-A and H5N8-B distribution and severity of lesions in different tissues of chickens

The histopathological changes and distribution of influenza antigen in different organs of chickens which died at 2 dpi were evaluated. All H5N8-A and H5N8-B viruses induced comparable levels of necrotic inflammation in different organs. Using IHC, virus antigen was detected in the endothelial cells and parenchyma of all organs at comparable levels (Figure 3C, D). These results revealed the systemic distribution of H5N8-A and H5N8-B viruses in chickens without obvious impact of NS1 CTE on lesions or distribution in vital organs.

### Elongation of NS1 of H5N8-B virus reduced virulence in Pekin ducks

Previous studies showed that H5N8-A virus was avirulent in Pekin ducks after ON inoculation, while H5N8-B was highly virulent (21). Furthermore, we showed that NS from H5N8-B virus increased virulence of H5N8-A in ducks (Scheibner et al. in preparation). Here, we assessed the impact of CTE on virus fitness in ducks. A_NS237 and the short NS1-derivatives exhibited low virulence in ducks and no mortality was observed (Figure 4A, B). All ducks seroconverted 10 dpi (data not shown). Conversely, all ducks inoculated with B_NS217 died within 4 days with MTD of 2.5 days (Figure 4C) and PI value of 2.7. Elongation of the NS1 CTE reduced H5N8-B virulence as indicated by reduced mortality and increased survival periods. After inoculation of ducks with B_NS230 and B_NS237, 9/10 and 7/10 ducks died with PI values of 1.8 and 1.7, respectively. The survival period was significantly longer; 6.5 and 4.4 days in ducks inoculated with B_NS230 (p < 0.0001) and B_NS237 (p < 0.001), respectively (Figure 4C). All co-housed ducks in groups inoculated with B_NS217 died within 6 dpi (5 dpc) with MTD of 4.6 dpi (3.6 dpc) (Figure 4D). Only 2/5 contact ducks died in either group inoculated with B_NS230 or B_NS237. The contact ducks died at 10 or 8 dpi (9 and 7 dpc), respectively which was significantly longer than contact ducks co-housed with B_NS217 (p =0.003) (Figure 4D). All surviving ducks seroconverted 10 dpi (data not shown). These data confirm that H5N8-B is more virulent than H5N8-A in Pekin ducks. NS1 alone does not play a main role in virulence or transmission of H5N8-A, but elongation of NS1 CTE reduced virulence of H5N8-B in inoculated and contact ducks.

**Figure 4:**
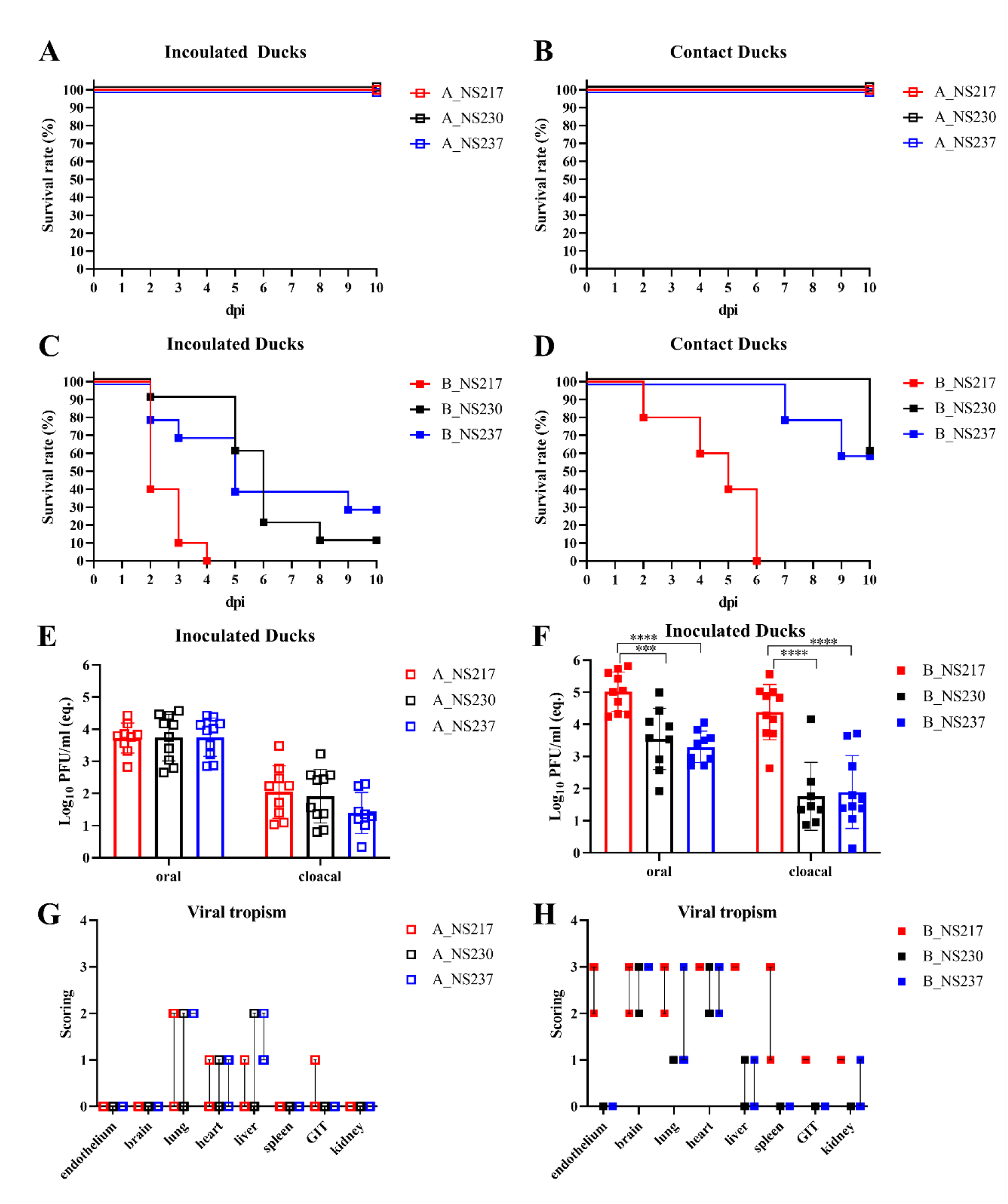
The impact of NS1 on virulence, transmission, virus excretion and endotheliotropism in ducks. Survival curves were generated using Kaplan Meyers for clade-A inoculated (A) and contact ducks (B) and clade-B inoculated (C) and contact ducks (D). Virus excretion determined in oropharyngeal and cloacal swabs in inoculated ducks with clade-A (E) or clade-B viruses (F) using quantitative RT-qPCR was expressed as log10 pfu/ml (eq.) in inoculated ducks. Viral tropism in ducks was determined by IHC at 2 dpi of ducks with clade-A (G) or clade-B (H) viruses. Scores indicate 0 = negative, 1 = focal to oligofocal, 2 = multifocal, 3 = coalescing foci or diffuse labelling, dots represent individual animals, bar = median with interquartile range. Endothelium scores indicate the maximum score found in all tissue affected. Statistical significance * p < 0.05, ** p< 0.01, ***, p <0.001, ****, p <0.0001.

### Changes in the NS1 CTE reduced H5N8-B virus excretion in cloacal swabs in ducks

Virus excretion was determined in OP and CL swabs obtained at 2 dpi in inoculated and 1 dpc in contact ducks. Inoculated ducks excreted more virus orally than via the cloacae. B_NS217 was excreted at significantly higher levels than A_NS237 in inoculated ducks (Figure 4E, F). No significant differences were observed in H5N8-A inoculated groups (Figure 4E), but the level of virus shedding in the OP swabs was significantly higher than in the CL swabs (p ≤ 0.0001). Conversely, B_NS217 was excreted at significantly higher levels than B_NS230 and B_NS217 (p < 0.005) with about 15 and 1000 average folds, respectively (Figure 4F). In contact ducks, all H5N8-A viruses were excreted at similarly low levels in the OP swabs and viral RNA was detected in 1/5 duck only in the cloacal swabs. B_NS217 and B_NS237 were detected at similar levels in the OP swabs, and at ∼10 times higher levels than B_NS230 (data not shown). Only 1/5 ducks cohoused with B_NS217 inoculated ducks excreted virus in the CL swabs (10^4^ pfu/ml), other contact ducks were negative (data not shown).

### Changes in the CTE altered H5N8-B distribution and lesions in different tissues of ducks

H5N8-A viruses induced focal to multifocal necrosis and inflammation in the lungs, brain, heart and nasal conchae, without impact of CTE on the distribution of the lesions. H5N8-B viruses caused more consistent and more widespread (multifocal to diffuse) necrosis and inflammation in these tissues and additionally in the spleen, thymus, bursa, pancreas and kidney. Remarkably, hepatic necrosis was decreased or abolished by gradually shortening the H5N8-A NS1 or extension of H5N8-B NS1. Viral antigen detection in the endothelium was only seen in B_NS217 with a broad spectrum of affected tissues (systemic endotheliotropism). In accordance with the lesion profile, H5N8-A viruses were mainly found in the lungs, heart, nasal conchae and liver but clearly less consistent and less abundant compared to H5N8-B viruses (Figure 4G,H). Interestingly, only A_NS237 showed viral antigen in the parabronchial epithelium of air sac ostia. The tropism of H5N8-B viruses was extended to the brain, thymus, spleen, intestine, pancreas and kidney but elongation of NS1 resulted in reduced M1 antigen detection in all organs. Neither B_NS230 nor B_NS237 was detected in the gastrointestinal tract (Figure 5, detailed data not shown). Together, compared to H5N8-A virus, the high virulence of H5N8-B virus in ducks was associated with more widespread necrosis and inflammation as well as lymphocyte depletion in the lymphatic organs and in case of B_NS217 with systemic endotheliotropism. The NS1 CTE abolished the diffuse endotheliotropism of H5N8-B virus, but alone did not affect the lack of endotheliotropism of H5N8-A virus and gradually reduced the hepatic necrosis caused by both H5N8-A and H5N8-B and the antigen distribution of H5N8-B in all tissues.

**Figure 5:**
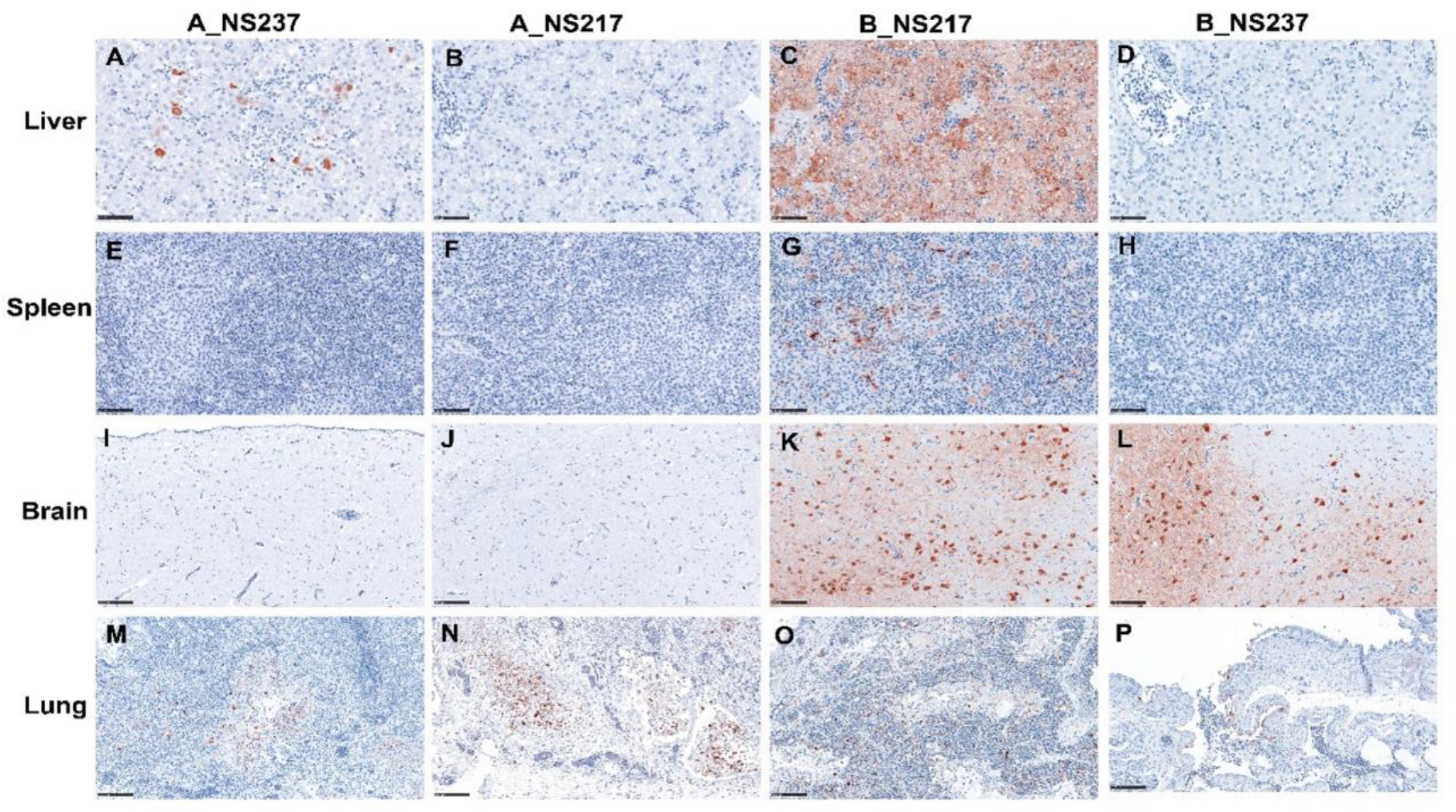
Immunohistopathological detection of Matrix antigen in selected organs in ducks. Organs were obtained from inoculated-ducks, euthanized 2 dpi. Immunohistochemistry, ABC Method using anti-Matrix-1 protein antibody, AEC chromogen (red-brown), Hematoxylin (blue) counterstain. (A-H) bar 50 µm, (I-P) bar 100 µm.

### Detection of cytokine response in the lungs and spleen of inoculated chickens and ducks

The detection of IFN-α, IFN-β and IFN-γ mRNA in the lungs and spleen of inoculated chickens and ducks at 2 dpi was done using generic RT-PCR and expression levels were normalized to 28S rRNA. The results are expressed as fold change compared to negative controls (Figure 6). In the lungs of chickens, H5N8-A and H5N8-B viruses induced comparable expression levels for IFN-α and IFN-γ, while IFN-β expression induced by A_NS237 (Figure 6B) was significantly lower than that induced by clade B_NS217 (p < 0.05) (Figure 6E). A_NS237 was more efficient to block IFN-α induction than A_NS230 and A_NS217 (Figure 6A) and was able to significantly block IFN-γ induction compared to A_NS230 Figure 6C). B_NS230 was significantly less efficient to inhibit IFN-α induction than B_NS217 and B_NS237 (Figure 6D). In the spleen, A_NS217 was less efficient than A_NS230 and A_NS237 in inhibiting the IFN-α response. No significant differences were observed in the spleen of chickens inoculated with H5N8-B viruses (data not shown). In ducks, the expression of IFN was limited compared to chickens. There were no significant differences in the levels of expression of IFN in the lungs and spleen between different groups (Figure 6G-L, data not shown). These results indicate that in chickens the original NS1 of H5N8-A and H5N8-B viruses, regardless of the length of the CTE, evolved toward higher efficiency to block IFN-α response in the lungs. Extension or shortening of the NS1 reduced the efficiency of the virus to block IFN response. IFN response in ducks was limited compared to chickens and NS1 CTE did not affect IFN expression in the lungs and spleen.

**Figure 6:**
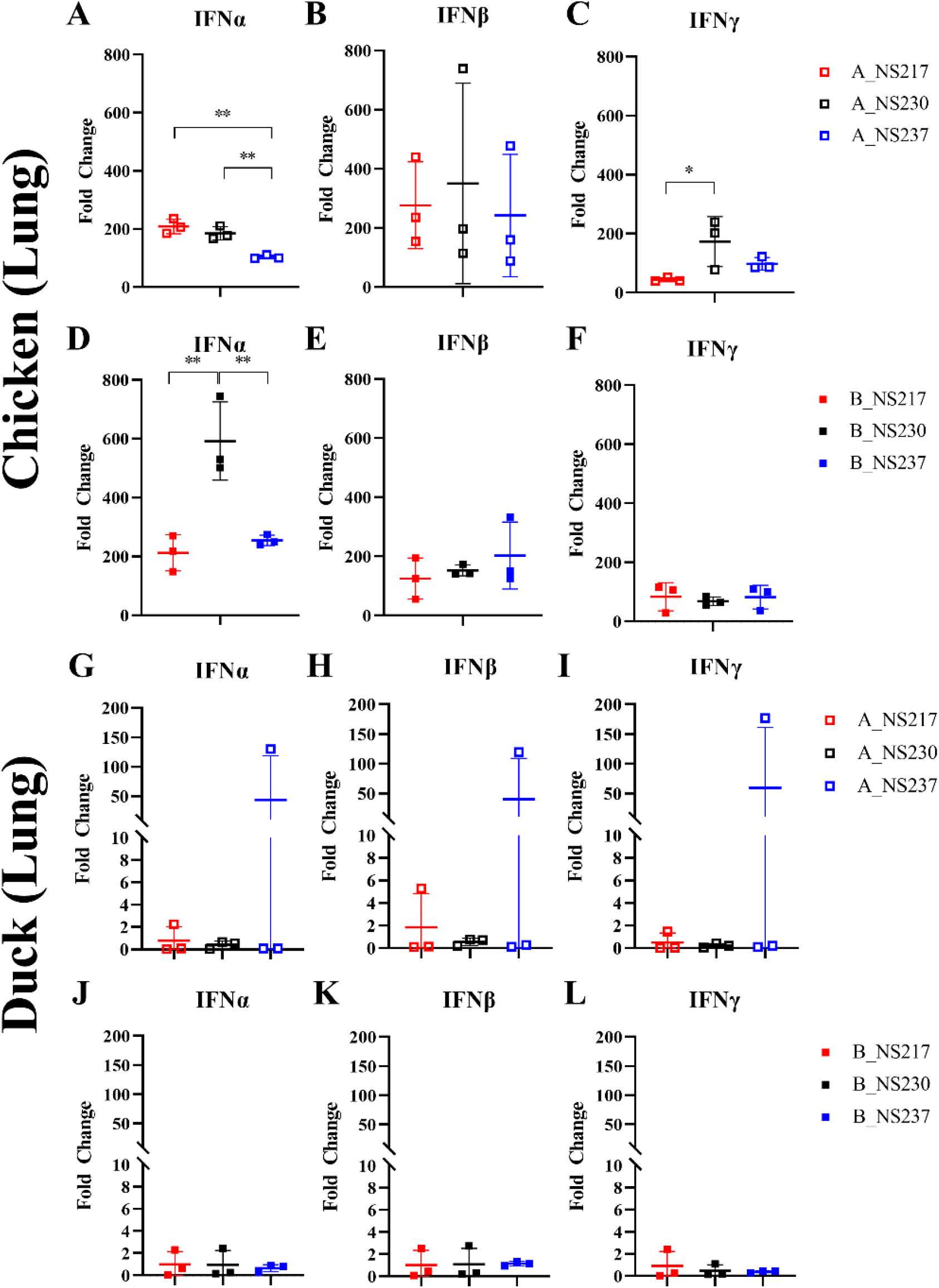
Interferon induction in lungs of chickens and ducks inoculated with clade A and B H5N8 viruses. Shown are the fold changes in IFN-levels in the lungs of chickens and ducks inoculated with clade A and B H5N8 viruses 2 dpi. Tissues were collected, weighed (w/v) and homogenized. The mRNA of IFN-α, IFN-β or IFN-γ were measured by generic RT-qPCR from 3 birds in each group. Normalization was done using 28S rRNA transcripts. Results were calculated using the 2^-(ΔΔct) method and expressed as fold change of normalized samples compared to samples ossbtained from non-infected birds (n=3). Statistical significance * p < 0.05, ** p< 0.01.

## Discussion

The continuous circulation of the panzootic H5N8 clade 2.3.4.4 is threatening the poultry industry worldwide. It is important to understand the viral factors which contribute to the high fitness of HPAIV H5N8 in chickens and ducks. It has been previously suggested that H5N8 clade-A viruses acquired the NS segment from A/duck/Eastern China/1111/2011 (H5N2), while clade-B viruses acquired the NS segment by reassortment with A/environment/Jiangxi/28/2009 (H11N9) or A/duck/Hunan/8-19/2009 (H4N2) (34). We found that all putative ancestors for clades A or B possess NS1 with 230-aa. In contrast, Eurasian H5N8 in 2013/2014 (clade-A) or European 2015/2016 (clade-B) have NS1 with 237-aa or 217-aa, respectively. We did not find NS217 in 2013/2014 or NS237 in 2016/2018. These results indicate that NS1 rapidly acquired longer (clade-A) or shorter (clade-B) NS1 and dominated their ancestors indicating selective advantages for virus replication in a clade-specific pattern. Indeed, our competition experiments in cell culture confirmed this assumption with preferential selection of the authentic NS over NS with a typical length of 230-aa. The latter was disadvantageous for virus fitness *in vitro* and/or *in vivo*. The negative impact of NS230 on the fitness of clade-A virus was limited compared to clade-B virus. In clade-A virus, the only significant difference due to shortening of the NS237 to 230-aa or 217-aa was the less ability to block IFN-α response in the lungs of chickens compared to the wild type A_NS237 virus. Conversely, the extension of NS1 CTE of clade-B virus to 230-aa or 237-aa reduced virus replication in duck cells and virulence, transmission, excretion and/or replication in different organs of inoculated chickens and ducks. These results may explain the preferential selection of H5N8 clade 2.3.4.4 for a certain length of the NS1 protein.

While chickens died after inoculation with H5N8-A and H5N8-B viruses, H5N8-B was more virulent in Pekin ducks than H5N8-A virus which is in accordance with previous results (21). The virulence determinants of HPAIV vary in chickens and ducks. In chickens, the HA, particularly the polybasic cleavage site (HACS), is the main determinant of virulence. However, mutations in other gene segments (i.e. PB1, NP and NA) contributed to the high virulence of an American HPAIV H5N2 clade 2.3.4.4A in chickens (35). Moreover, previous studies have shown that NS1 V149A contributed to the virulence of an H5N1 virus in chickens (36) and replacing the NS segment of the current H5N8-B with that of an H9N2 reduced transmission and replication of H5N8-B in chickens (22). In ducks, several studies found that the HA alone is not sufficient for high virulence (37-42) and we recently found that H5N8-B NS segment, in addition to the HA and NP, increased virulence and transmission of H5N8-A in Pekin ducks (Scheibner et al. in preparation) indicating a significant role for NS in H5N8-B virulence and bird-to-bird transmission efficiency. Here, we showed that the NS1 CTE is important for virus transmission in chickens and ducks in a virus-specific manner. Although it remains to be studied, H5N8-A virus may have compensatory mutations in the NS1 or other proteins and, thus, shortening the CTE have less impact on virus fitness compared to H5N8-B. In fact, the synergism between NS1 CTE and mutations in the NS1 RBD (i.e. I38Y) (43) or mutations in the PB1-F2 (44) affected virulence of HPAIV H5N1 in mice. Similarly, mutations in the nuclear export signal (e.g. D139N as seen in this study) can compensate the absence of nuclear/nucleolus localisation signals due to a 6-aa-deletion in the NS1 CTE of an H7N7 virus (12).

Virulence in inoculated chickens and ducks was associated with systemic dissemination of viral antigen, histopathological lesions and in particular with systemic endotheliotropism in chickens and H5N8-B virus in ducks. Interestingly, in contrast to the systemic replication of H5N8-A virus in chickens, replication of H5N8-A virus in ducks was limited to the respiratory tract, heart and liver resembling some LPAIV and HPAIV (45). Viral or host factors which contribute to the different endotheliotropism in chickens and ducks of some AIV are not well understood (46). It has been shown that the polybasic HACS is important for the endotheliotropism in chickens and/or ducks (47, 48). However, although both H5N8 viruses used in this study possess a polybasic HACS, they showed a striking difference in endotheliotropism in ducks indicating that factors beyond the HACS are essential for the tropism to endothelial cells. Scheibner et al. (49) described diffuse endotheliotropism in chickens but not in ducks after inoculation with an HPAIV H7N7. In this study, the NS1 CTE did not significantly affect the endotheliotropism in chickens, however, it reduced H5N8-B virus distribution in the endothelial cells as well as in vital organs including the brain, heart, lung and spleen in ducks. The extension of CTE in HPAIV H7N1 NS1 decreased virus excretion and tropism to the endothelial cells and epithelium in the central nervous system and respiratory tract without significant difference in virulence in chickens (8). The specific impact of NS1 CTE on the endotheliotropism of H5N8-B virus in ducks merits further investigation.

Chickens mount robust IFN-responses, but fail to limit viral replication and succumb to a “cytokine storm” (50). Compared to chickens, we found that the IFN response in the lungs and spleen in Pekin ducks was limited as described before (51-55). Interestingly, NS1 is a main antiviral antagonist for influenza viruses which is mediated by different NS1 domains (6). Our results showed that CTE has no impact on IFN-β and IFN-γ responses, while H5N8-A and H5N8-B viruses with NS of 230-aa were less efficient to block IFN-α induction in chickens which might also explain the disfavor to NS with 230-aa. A previous study has shown that extension of NS1 of H9N2 to 230-aa or 237-aa did not affect the levels of IFN-α and IFN-β but increased the IFN-γ in the lungs of chickens (13). Conversely, a deletion of 6-aa in the NS1 CTE of an LPAIV H7N1 did not affect type I or type II IFN-response in chickens or ducks (11). This discrepancy is probably due to the use of different virus strains or subtypes.

In conclusion, there is a preferential selection for a certain NS1 CTE in 2.3.4.4 H5N8 clade-A (with 237-aa) and clade-B (with 217-aa) viruses over NS1 with 230-aa, the common length of NS1 in AIV. The latter had a negative impact on virus fitness *in vitro* and *in vivo*. CTE can affect the virulence in a species and clade-specific manners. In chickens, NS1 CTE of H5N8-A and H5N8-B evolved toward higher efficiency to block IFN-α response. In ducks, NS1 CTE is essential for efficient transmission, replication and high virulence of H5N8-B which correlated with (i) systemic endotheliotropism and (ii) widespread tissue damage. These results are important to understand the evolution of the panzootic H5N8 clade 2.3.4.4 and the role of NS1 in virus fitness in chickens and ducks.

## Acknowledgment

Dajana Helke, Nadine Bock, Sarah Knapp and Silvia Schuparis are thanked for laboratory technical assistance, Günter Strebelow for his assistance in sequencing, Dr. Christine Fast, Prof. Steffen Weigend, Bärbel Hammerschmidt, Frank Klipp, Harald Manthei, Doreen Fielder, Bärbel Berger, Thomas Moeritz and Ralf Henkel for their support in the animal experiments, Silvia Schuparis for histological preparations and Dr. Daniel Marc for providing the anti-NS1 antibodies. Emmelie Eckhardt and Luca Zaeck are thanked for technical support. This work was supported by grants from Horizon 2020 “Delta Flu” (project ID: 727922) and the Deutsche Forschungsgemeinschaft (DFG; AB 567/1-1 and DFG VE780/1-1). The funders had no role in study design, data collection and analysis, decision to publish, or preparation of the manuscript.

## Declaration of interest statement

The authors declare no conflict of interest

